# Resistance to host antimicrobial peptides mediates resilience of gut commensals during infection and aging in *Drosophila*

**DOI:** 10.1101/2023.03.21.533593

**Authors:** Aranzazu Arias-Rojas, Dagmar Frahm, Robert Hurwitz, Volker Brinkmann, Igor Iatsenko

## Abstract

Resilience to short-term perturbations, like inflammation, is a fundamental feature of microbiota, yet the underlying mechanisms of microbiota resilience are incompletely understood. Here we show that *Lactiplantibacillus plantarum*, a major *Drosophila* commensal, stably colonizes the fruit fly gut during infection and is resistant to *Drosophila* antimicrobial peptides (AMPs). By transposon screening, we identified *L. plantarum* mutants sensitive to AMPs. These mutants were impaired in peptidoglycan O-acetylation or teichoic acid D-alanylation, resulting in increased negative cell surface charge and higher affinity to cationic AMPs. AMP-sensitive mutants were cleared from the gut after infection and aging-induced gut inflammation in wild-type, but not in AMP-deficient flies, suggesting that resistance to host AMPs is essential for commensal resilience in an inflamed gut environment. Thus, our work reveals that in addition to the host immune tolerance to the microbiota, commensal-encoded resilience mechanisms are necessary to maintain the stable association between host and microbiota during inflammation.

## Introduction

Gut-associated microbial communities stably colonize the host over the lifetime of an individual despite constant exposure to perturbations which often transiently change microbial community function and composition^1^. Over time, the community can revert to the original state as the disturbance passes, thus exhibiting resilience^2, 3^. However, when resilience mechanisms fail, perturbations may lead to the establishment of dysbiosis, a disease-associated state of the microbiota, which negatively influences the health of the host^4^. Hence, understanding the molecular mechanisms of microbiota resilience is important for maintaining the health of the host. Some perturbations to which the intestinal microbiota is often exposed to are non-specific inflammatory responses induced by infection. Immune defence mechanisms non-specifically target conserved molecular patterns present in both pathogens and commensals, raising the question of how commensals survive such inflammatory responses and can stably colonize the host. Recent studies have started addressing this question. For instance, human commensal Bacteroidetes modify their lipopolysaccharide (LPS) structure, leading to increased resistance to host antimicrobial peptides and persistence in the gut during inflammation^5^. Another perturbation to which commensals are exposed to during infection is an inflammation-induced iron limitation. *Bacteroides thetaiotaomicron* survives such iron-limiting conditions by utilizing siderophores produced by members of the *Enterobacteriaceae* family to acquire iron^6^. This xenosiderophore utilization suggests a crucial role for interspecies iron metabolism in mediating commensal resilience during gut inflammation. However, despite the latest efforts, our understanding of the molecular mechanisms that underlie microbiota resilience during infection remains limited.

One approach to explore microbiota resilience mechanisms towards immune defences is to use *in vivo* model systems amenable to genetic manipulation of both host and microbiota members. The fruit fly *Drosophila melanogaster* is one such model that has been widely used to study host-microbe interactions^7–9^.

Similar to the mammalian intestinal tract, the *Drosophila* gut is equipped with barriers that control bacterial proliferation and prevent microbe-induced damage to the gut epithelia^10, 11^. Specifically, ingested pathogens induce two types of effectors that act synergistically to restrict the growth of intestinal microorganisms: antimicrobial peptides and reactive oxygen species (ROS)^12–14^. Infection-induced expression of AMPs in specific regions of the gut is the hallmark of the *Drosophila* immune response^15^. The AMP response is regulated by two conserved nuclear factor-kB (NF-κB) pathways: Toll and immune deficiency (Imd)^16, 17^. The Imd pathway is initiated in the gut when diaminopimelic (DAP)-type peptidoglycan from bacteria is sensed by the transmembrane recognition receptor PGRP-LC in the ectodermal parts of the gut or by the intracellular receptor PGRP-LE in the midgut, ultimately leading to the nuclear translocation of the NF-kB transcription factor Relish^18, 19^. Activated Relish then induces the expression of immune effectors, like AMPs, that eliminate the pathogens.

With the advent of microbiome research, it became apparent that Imd also responds to commensals and mediates their impact on several physiological processes. Indeed, the relatively simple microbiota of fruit flies, consisting of about 30 phylotypes and dominated by *Lactobacillaceae* and *Acetobacteraceae*, has been shown to have profound effects on intestinal metabolism, immune response, and tissue homeostasis^9, 20, 21^. Most of the *Drosophila* microbiota members, similar to pathogens, produce DAP-type PGN – a major elicitor of the Imd pathway. However, the microbiota remains only a mild inducer of the AMP response, suggesting that flies deploy immune tolerance mechanisms to commensals. Indeed, while commensals stimulate a weak AMP response, they induce a strong expression of negative regulators of the Imd pathway^19, 22, 23^. These negative regulators, like the PGN-degrading enzymes PGRP-SC1 and PGRP-LB, maintain a low basal level of immunostimulatory PGN, thus preventing the overactivation of the Imd pathway to the gut microbiota^24–26^. On one hand, these negative regulators protect the host from chronic deleterious Imd pathway activation, on the other hand, they prevent a strong AMP response that would target gut commensals. However, during infection this host-microbiota homeostasis is disrupted as pathogens trigger a transient but strong production of AMPs. These AMPs not only neutralize pathogens, but will also target gut commensals. Therefore, the question of how the composition and abundance of intestinal microbiota is affected by infection and how it tolerates the exposure to AMPs remains to be addressed. Similarly, during aging, flies exhibit increased commensal loads despite an elevated AMP response^27, 28^, raising the question of how commensals are able to survive in these inflamed environments.

Here, we showed that *Drosophila* microbiota composition and abundance remains stable during infection. Using the dominant *Drosophila* commensal *L. plantarum* as a model, we discovered that it does not avoid the immune response but is resistant to cationic AMPs. In a transposon screen, we identified several AMP-sensitive *L. plantarum* mutants with altered cell wall modifications and increased binding of AMPs. The abundance of AMP-sensitive mutants was significantly reduced after infection in wild-type but not in AMP-deficient flies. Taken together, our results demonstrate that the resistance to host AMPs via cell wall modifications is crucial for microbiota resilience during infection.

## Results

### *Drosophila* microbiota is resistant to the host intestinal immune responses induced by oral infection

First, we investigated the impact of infection on *Drosophila* microbiota composition using 16s rRNA sequencing of dissected gut samples of 10d old conventional (colonized with native microbiota) wild-type iso (*w^1118 iso^*) flies infected with alive and heat-killed *Erwinia carotovora (Ecc15)* (Fig. 1a). Heat-inactivated *Ecc15* induces the IMD pathway but does not directly interfere with commensals. Treatment of flies with *Ecc15*, either live or heat-killed, did not decrease the microbiota species richness and diversity as illustrated by Shannon and Simpson indexes (Fig. 1b, 1c, Supplementary Fig. 1a-c) but shifted dissimilarities in the treatments over time (Fig. 1d), suggesting that some members of the microbiota can tolerate the intestinal immune response better than others. To determine which microbiota members are resistant to the immune response, we analysed the relative abundance of the most representative species in our 16S sequencing data. Consistent with previous studies^29, 30^, microbial communities in our flies were dominated by the phylum of Firmicutes represented by the families of *Lactobacillaceae* and *Enterococcaceae* and the phylum of Proteobacteria represented by the families of *Sphingomonadaceae* and *Enterobacteriaceae* (Supplementary Fig. 1d). We noticed that infection led to a reduction in *Sphingomonadales* and a stable or even increased abundance of *Clostridiales*, and *Lactobacillales* species (Fig. 1e). Overall, while there were minor changes in the composition after infection, diversity indexes did not show any statistically significant changes in microbiota composition between uninfected and infected samples, suggesting that the *Drosophila* intestinal microbiota community is relatively stable and can persist infection-induced immune and stress responses in the gut.

**Figure 1.**
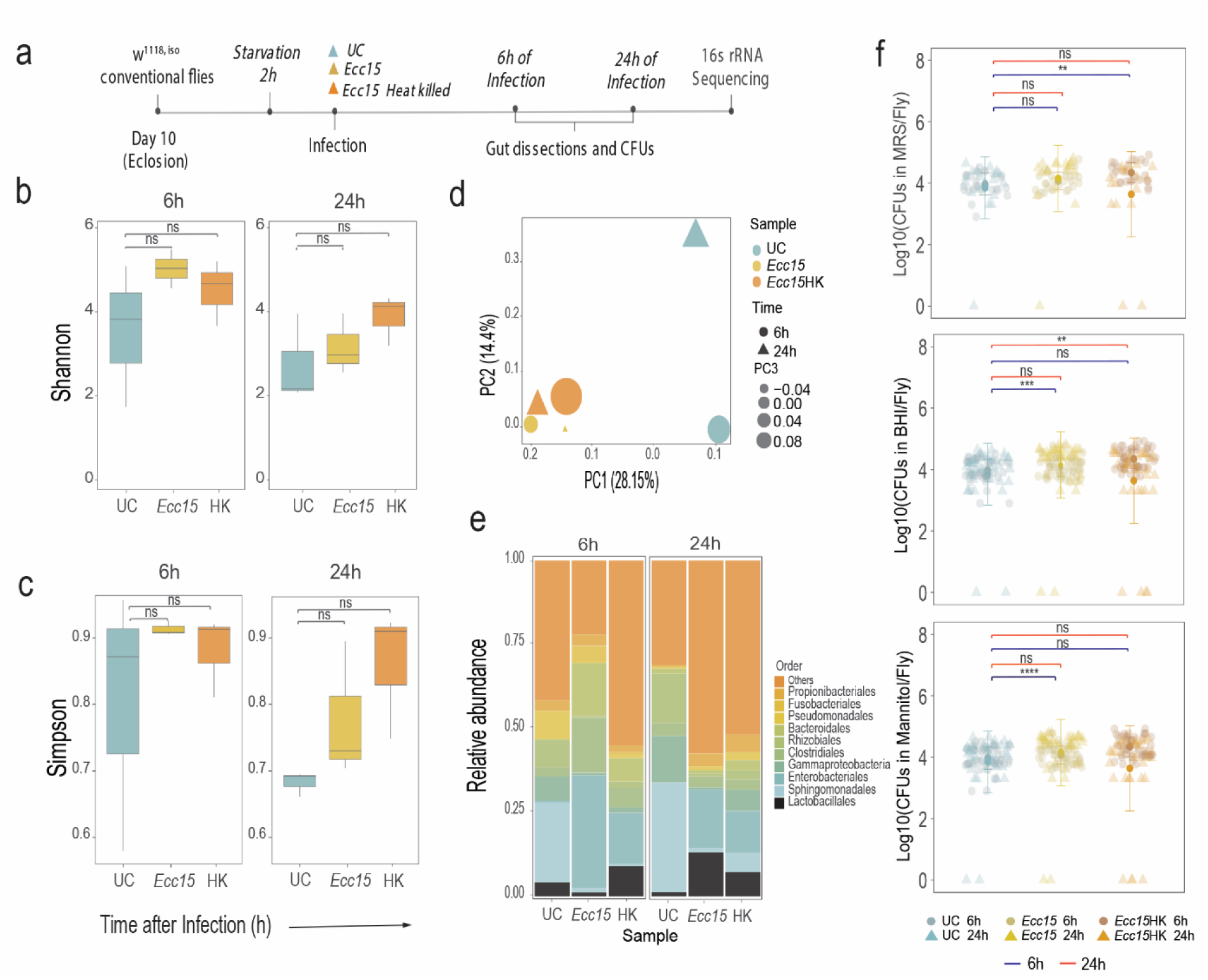
*Drosophila* microbiota is resistant to the host intestinal immune responses induced by oral infection. **a,** experimental design to investigate infection’s effect on microbiota composition and abundance in 10d old conventional flies. (Unless otherwise stated, n=3 independent experiments with 20 guts per treatment performed at 6h and 24h). **b-c**, alpha diversity illustrated by Shannon (**b**) and Simpson (**c**) indexes throughout the time. **d**, beta diversity, illustrated by unweighted unifrac distances among treatments and time points. **e**, the relative abundance of 10 dominant OTUs after *Ecc15* infection. **f**, culturable microbiota loads in 10d old conventionally reared flies 6h and 24h post *Ecc15* oral infection in MRS, BHI, and mannitol agar. Sample size: UC 6h (n=26), UC 24h (n=18), *Ecc15* 6h (n=26), *Ecc15* 24h (n=19), *Ecc15*-HK 6h (n=25*), Ecc15*-HK 24h (n=17). The single dots are mean CFU values from pools of n = 5 animals in the Log10 scale. Dot plots and boxplots show median and interquartile range (IQR), and whiskers show either the lower and upper quartiles or range. *P < 0.05, **P < 0.01, ***P < 0.001, ****P < 0.0001. Kruskal–Wallis and Bonferroni post hoc tests were used for statistical analysis.

Next, to corroborate our 16S rRNA sequencing results, we recorded the microbiota loads of whole 10d conventional flies infected with heat-killed *Ecc15* by plating fly homogenates on media supporting microbiota growth (MRS, Mannitol, and BHI). Treatment with *Ecc15* had no significant effect on microbiota load in most cases (Fig. 1f). In some cases, we even observed an increase in microbiota cell numbers after infection (Fig. 1f).

In the next set of experiments, we used gnotobiotic flies monocolonized with representative microbiota isolates *L. plantarum or A. melorum* and measured their persistence after infection with alive and heat-killed *Ecc15,* respectively (Supplementary Fig. 2a). We found that in most cases the loads of the tested species were not significantly affected by the infection (Fig. 2a, 2b), which we confirmed through qPCR. This was also true for the different *L. plantarum* strains that we tested (Supplementary Fig. 2b-e). Consistent with our 16S sequencing data (Fig. 1e), the numbers of *L. plantarum* were even increased 24h after *Ecc15* infection (Fig. 2a and Supplementary Fig. 2b-d). Similarly, we detected a significantly elevated amount of *A. melorum* cells 24h post infection (Fig. 2b). Taken together, these results show that representative members of the *Drosophila* microbiota are able to persist in the gut during intestinal immune response and some of them even increase in numbers.

**Figure 2.**
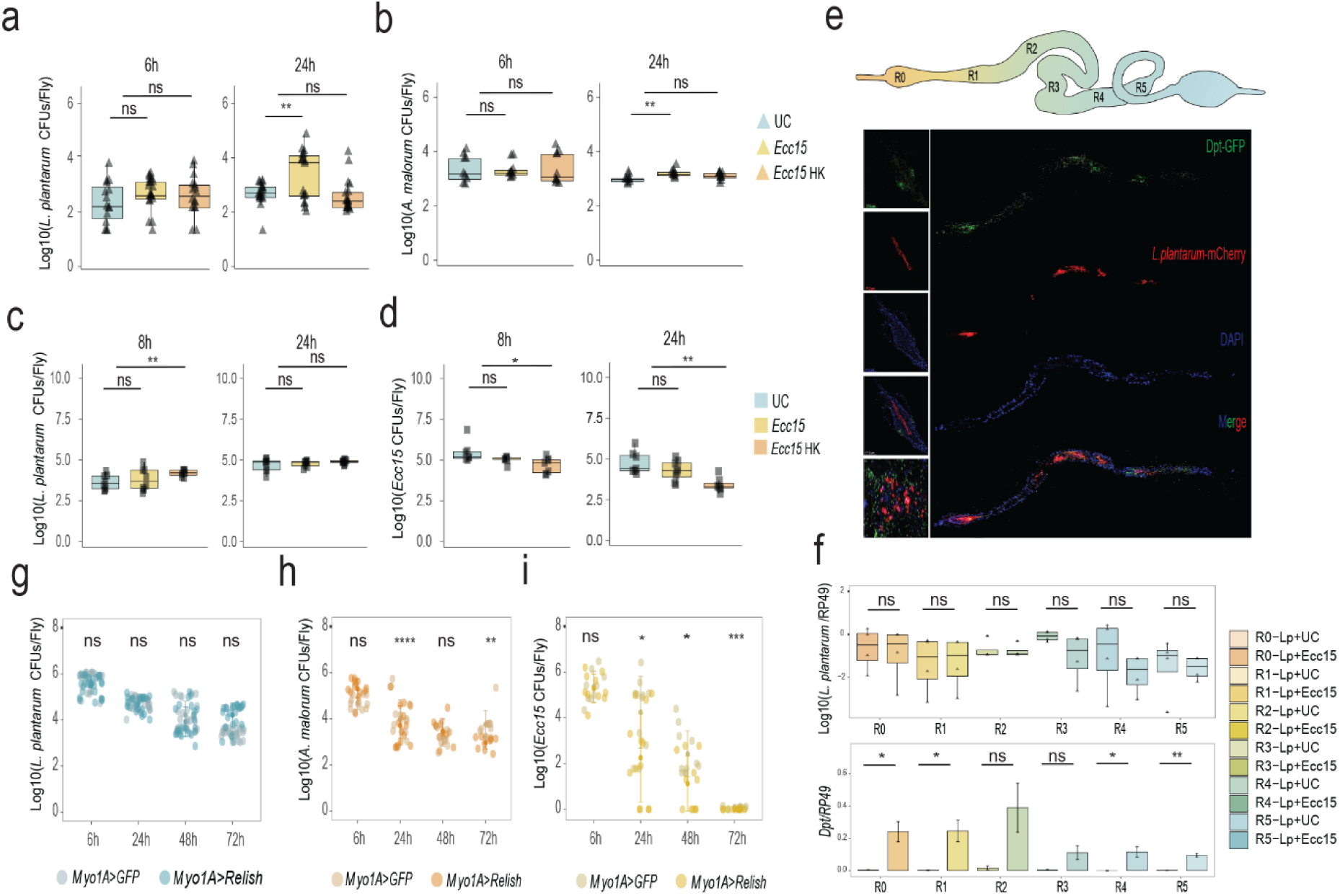
*L. plantarum* is present in the immune-responsive gut regions and is resistant to the IMD pathway effectors. **a-b**, *L. plantarum^NCIMB^* (**a**) and *A. malorum* (**b**) loads 6h and 24h after the infection with *Ecc15*. **c-d**, *L. plantarum* (**c**) and *Ecc15* (**d**) loads 8h and 24h in flies that were orally primed with *Ecc15* before colonization (n=8 independent samples per treatment with 5 flies per sample). The single dots are mean CFU values from pools of n = 5 animals in the Log10 scale. **e**, Imaging of adult DptB-GFP *Drosophila* gut colonized with *L. plantarum*-mCherry^WJL^ and infected with *Ecc15* after 6h. **f**, *L. plantarum^WJL^* loads recorded by qPCR 6h after *Ecc15* infection, matched with *DptA* expression per gut region in *w^1118^* iso flies. Bar plots show media and se range (n=3 independent experiments). **g**, CFUs of *L. plantarum^NCIMB^* in *Myo1A-GAL4>UAS-GFP* and *Myo-GAL4>UAS-Relish* flies at 6h, 24h, 48h, and 72h post colonization (n=24 for both genotypes at all time points). **h**, CFUs of *A. malorum* at 6h (n1=12 and, n2=13), 24h (n1= 14 and, n2=14), 48h (n1=11 and, n2=10), and 72h (n1=10 and, n2=10) post colonization in *Myo1A-GAL4>UAS-GFP* (n1) and *Myo-GAL4>UAS-Relish* flies (n2). **i,** *Ecc15* loads at 6h (n1=9 and, n2=10), 24h (n1= 11 and, n2=13), 48h (n1=11 and, n2=11), and 72h (n1=10 and, n2=10) post infection in *Myo1A-GAL4>UAS-GFP* (n1) and *Myo-GAL4>UAS-Relish* flies (n2). The single dots are mean CFU values from pools of n = 5 animals in the Log10 scale. Boxplots and dot plots show median and interquartile ranges (IQR); whiskers show either lower or upper quartiles or ranges. *P < 0.05, **P < 0.01, ***P < 0.001, ****P < 0.0001. Kruskal–Wallis and Bonferroni post hoc tests were used for statistical analysis.

Additionally, we investigated whether intestinal immune activation prior to colonization (priming) will affect the ability of representative commensals to colonize and persist in the gut. To address the effect of immune priming, we infected wild-type germ-free flies with either alive or heat-killed *Ecc15* and performed colonization with certain microbiota members (Supplementary Fig. 2a). *L. plantarum* showed an increase in the *Ecc15* heat-killed treatment at 8h and no significant changes under the other conditions (Fig. 2c). We confirmed these results by qPCR, and for different *L. plantarum* strains (Supplementary Fig. 2f-i). Importantly, when we infected *Ecc15*- primed flies with *Ecc15* or *P. entomophila (Pe)*, we observed reduced *Ecc15* or *Pe* loads compared to control flies (Fig. 2d and Supplementary Fig. 2j-k). This result proves that priming (1) induces an immune response and (2) that this immune response is effective against pathogens but not the microbiota members that we tested.

Altogether, our results demonstrate that infection causes only minor changes in the composition of the *Drosophila* microbiota and that the dominant gut commensals can persist in the gut during active immune responses induced by pathogens.

### *L. plantarum* is present in the immune-responsive gut regions

Next, we focused on one of the dominant commensals in our flies, *L. plantarum*, to understand how it persists in the gut during infection. One hypothesis we had is that *L. plantarum* might simply avoid regions where the immune system is active and thereby survive the immune challenge. This scenario is possible, considering that the *Drosophila* gut is regionalized (Fig. 2e) and that the gut regions differ in the intensity of the immune response^31^. We took several approaches to investigate this avoidance hypothesis. First, we infected flies that had been monocolonized with *L. plantarum* with *Ecc15* and dissected guts 6h post infection. The dissected guts were separated into 6 regions. We then performed qPCR to quantify *L. plantarum* and measure activation of the Imd pathway in the same intestinal region.

As shown in Fig. 2f, *Ecc15* infection potently induced *Diptericin A* expression (Imd pathway readout) in most of the gut regions. *L. plantarum* was detected in all the gut regions and its abundance was not affected by infection. Consistently, we also did not find increased *L. plantarum* loads in the gut regions lacking *DptA* induction after infection. Overall, this suggests that *L. plantarum* colonizes all *Drosophila* gut regions, regardless of the level of their active immune response. Additionally, we used a microscopy approach to visualize both immune activation and *L. plantarum* localization in the gut. Specifically, we colonized *DptB-GFP* reporter flies with *L. plantarum-mCherry* bacteria and visualized the localization of both 6h after *Ecc15* infection. We observed (Fig. 2e) an abundant localization of *Lp-mCherry* in the foregut region where *Dpt-GFP* expression was also very pronounced. A similar colocalization pattern was also seen in the other gut regions, confirming that *L. plantarum* does not avoid regions with active immune responses. Finally, we genetically overactivated the Imd pathway in enterocytes, a major intestinal cell type, by overexpressing the transcription factor Relish. While this intestine-wide immune activation significantly reduced the colonization of the gut by *Ecc15* (Fig. 2i and Supplementary Fig. 2m), which validated our assay, it did not significantly affect the abundance of *L. plantarum* (Fig. 2g and Supplementary Fig. 2l) or *A. melorum* (Fig. 2h) across several timepoints. Therefore, it is likely that gut commensals have a mechanism other than avoidance to withstand the action of intestinal immune defences.

### *Drosophila* commensals are resistant to cationic AMPs *in vitro*

We hypothesized that the *Drosophila* microbiota members are resistant to host antimicrobial peptides and, therefore, can survive infection-induced immune responses. Since it is not possible to obtain *in vitro* the exact combination of AMPs that is produced by intestinal cells of the fruit fly, we decided to test the readily available cationic antimicrobial peptide polymyxin B. It has been widely used to model AMP sensitivity and mimics the action of some *Drosophila* AMPs^5, 32, 33^. MIC test showed that typical *Drosophila* gut commensals, like *L. plantarum*, *L. brevis*, *A. melorum,* and *E. faecalis* are resistant to polymyxin B and still grow at the highest tested concentrations. In contrast, oral pathogens of fruit flies, including *Ecc15*, *P. entomophila,* and *P. aeruginosa* were sensitive and did not grow even at the lowest concentration of polymyxin B (Fig. 3a). Additionally, we estimated the proportion of polymyxin resistant microbes in our lab flies by plating fly homogenates on growth media supplemented with or without the antibiotic. While polymyxin supplementation had no effect on the number of bacteria growing on MRS and BHI, we observed a slight reduction in the number of mannitol-growing bacteria in the presence of the antibiotic. This indicates that the majority of culturable microbes in our flies are resistant to polymyxin B (Fig. 3b) and thus likely also to some of the *Drosophila* AMPs.

**Figure 3.**
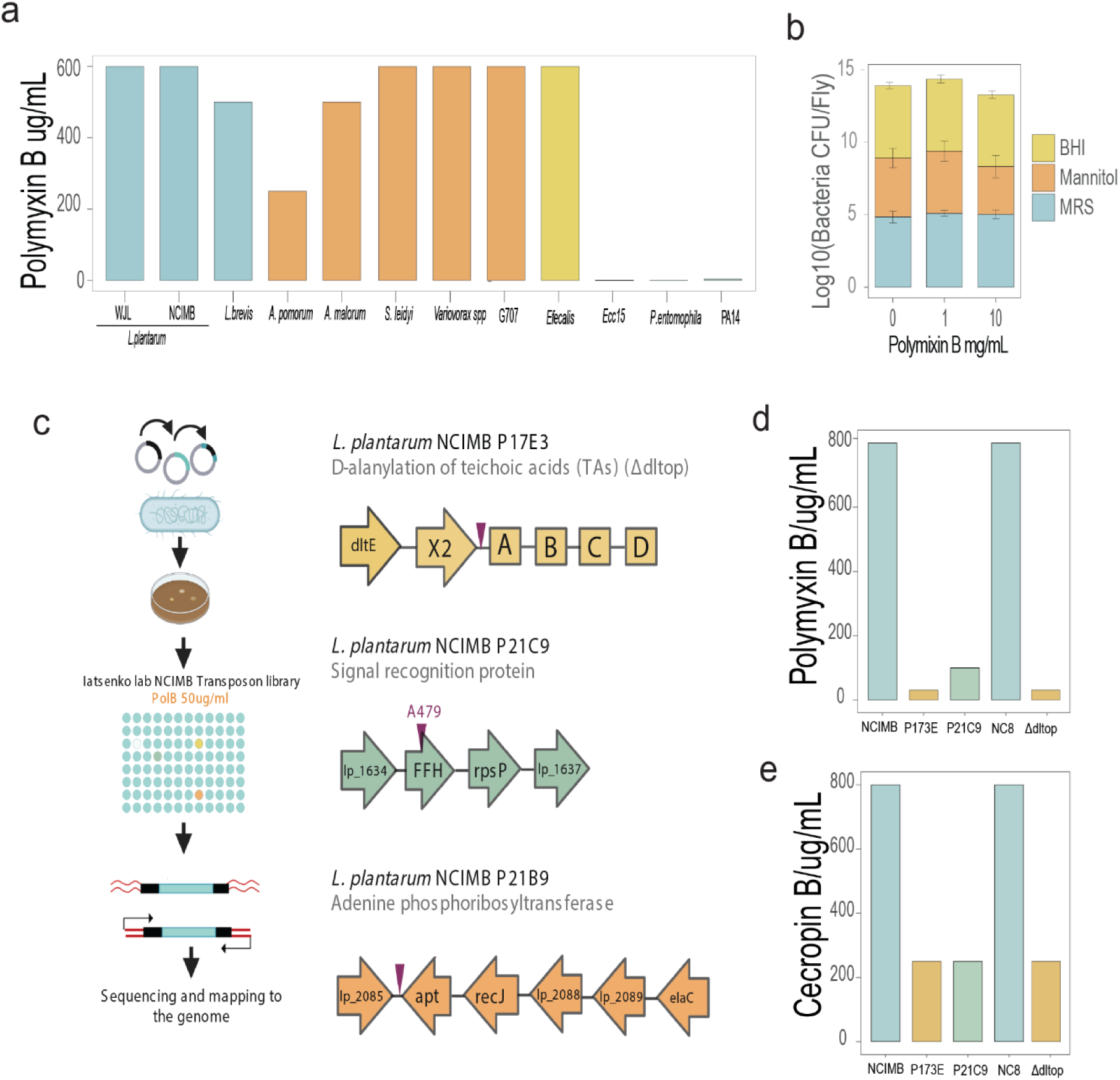
*Drosophila* microbiota is resistant to the AMPs *in vitro*. **a**, Minimal Inhibitory Concentration test (MIC) of *Drosophila* microbiota members and pathogens sensitivity to Polymyxin B in broth. **b**, Amount of culturable microbes derived from 10d old flies able to grow on MRS, Mannitol, and BHI agar plates supplemented with Polymyxin B. **c**, Scheme of the transposon screening in *L. plantarum^NCIMB^* for polymyxin B-sensitive mutants and location of transposon insertions. Genomic organization of *dlt* operon in *L. plantarum^NCIMB^*, insertion between the *dltX* and *dltA* genes; lp_1634 operon, where the insertion hits at the end of *ffh* gene, and *lp_2085* where the insertion hits in between the *lp_2085* operon and *apt*. **d-c**, MIC of *L. plantarum^NCIMB^,* P17E3, P21C9 transposon mutants and *L. plantarum^NC8^* and Δ*dltop* mutant in broth supplemented with either Polymyxin B (**d**) or Cecropin B (**e**). Bar plots show mean values. MIC experiments were repeated three times with identical results, therefore no error bars are shown.

### Identification of AMP-sensitive *L. plantarum* mutants

Next, we investigated the genetic bases of this resistance in *L. plantarum*. We took a classical unbiased forward genetic approach in which a library of bacterial mutants was screened for sensitivity to polymyxin B. First, we generated a random transposon mutant library in *L. plantarum* NCIMB8826 strain, which has a high transformation efficiency in contrast to *Drosophila* isolates. We mutagenized *L. plantarum* NCIMB8826 using the previously described P_junc_-TpaseIS *_1223_* transposon mutagenesis system^34, 35^ and randomly selected and stocked 3000 colonies as individual clones at −80 °C. We screened this library for mutants that were unable to grow in the presence of 50 µg/ml of polymyxin B, which is at least 10 times lower than the wild-type *L. plantarum* could survive (Fig. 3c). Under these selective conditions, initially we identified 15 potential candidates. However, we could not confirm the sensitivity for 6 of them in the second test, and another 6 candidates exhibited impaired growth even in the absence of polymyxin B. Therefore, only 3 mutants passed the selection criteria. In one of these mutants (P21B9), the transposon insertion happened in an intergenic region between the ORFs encoding adenine phosphoribosyltransferase (*apt*) and a transcription regulator (*lp_2085*) (Fig. 3c). Given that such an intergenic insertion can affect either an upstream or downstream ORF, or both, we decided not to consider this complex case in this study. In mutant P21C9, the transposon hit the *ffh* (fifty-four homologue) gene which is part of the signal recognition particle pathway (SRP) necessary for protein translocation, secretion, and membrane incorporation^36^. In the P17E3 mutant, the transposon was inserted between the *dltX* and *dltA* genes of the *dlt* operon (Fig. 3c). The *dlt* operon is responsible for the esterification of wall teichoic acids (WTAs) with d-alanine, thus reducing their negative charge and attraction of cationic AMPs to the bacterial cell wall^37, 38^. Consequently, mutants of the *dlt* operon are susceptible to AMPs, which we confirmed here for *L. plantarum*. Using a MIC test, we verified that P17E3 and P21C9 are indeed several times more sensitive to polymyxin B than wild-type bacteria (Fig. 3d). Interestingly, our P17E3 mutant exhibited the same level of sensitivity as an *L. plantarum* mutant lacking the entire *dlt* operon (*Δdltop)* (Fig. 3d-e), suggesting that the entire operon in P17E3 is likely disrupted. P17E3, *L. plantarum Δdltop*, and P21C9 mutants were also more susceptible to the *Drosophila* AMP cecropin B (Fig. 3e), confirming that our *L. plantarum* mutants are sensitive to other cationic AMPs and not specifically to polymyxin B. We decided to further characterize P21C9 (*ffh*) and P17E3 (*dltop*) mutants and investigate the reason for their sensitivity to AMPs and its consequences for the interactions with the host.

### Disruption of the *dlt* operon and the *ffh* gene increases binding of cationic AMPs to the cell surface

Considering that the cell surface is a key mediator of AMP-bacteria interactions, we used SEM to explore cell morphology and measure cell parameters of the P17E3 and P21C9 mutants. We did not observe any obvious morphological alternations in both mutants (Fig. 4a), with the exception that their cells appeared smaller compared to wild-type cells. Quantification of cell length confirmed that P17E3 and P21C9 cells are significantly shorter than wild-type cells (Fig. 4b). Consistent with a previous study^39^, we detected a significant reduction in the width of P17E3 cells (Fig. 4c), while P21C9 cells were slightly wider compared to wild-type cells Fig. 4c. Next, we investigated whether the observed morphological alternations affected the cell surface properties of the P21C9 and P17E3 mutants. Both mutants bound more cationic cytochrome C compared to wild-type bacteria, indicating an increased negative surface charge (Fig. 4d). Consequently, binding of the labelled cationic AMP 5-FAM-LC-LL37 was increased to P21C9 and P17E3 cells (Fig. 4e). Using HPLC, we confirmed that P17E3 cells, consistent with the function of the *dlt* operon, have reduced levels of D-alanine esterification of WTAs (Fig. 4f). In the P21C9 mutant, we detected an amount of D-ala released from WTAs that was comparable to wild-type bacteria (Fig. 4f), suggesting that a different cell wall modification affects the surface charge in this mutant. Considering that the disrupted *ffh* gene in the P21C9 mutant is part of the signal recognition particle pathway necessary for protein translocation and secretion, we hypothesized that the P21C9 mutant might be impaired in the secretion of certain proteins necessary for cell wall biogenesis or modifications as previously reported in different bacteria^40–42^. To test this hypothesis, we performed a proteomic analysis of secreted and surface-associated proteins in the P21C9 mutant and wild-type bacteria. In total, we detected 111 proteins, however only 25 proteins passed p-value and fold change significance cutoffs (Supplementary table 1). Most proteins (21 out of 25) showed a reduced abundance in the P21C9 mutant, which would be consistent with the theory of impaired protein secretion. Among proteins with reduced abundance, two acyltransferases (OatA (lp_0856) and OatB (lp_0925)) implicated in PGN O-acetylation^43^ caught our attention (Fig. 4g). Since PGN O-acetylation mediates sensitivity to lysozyme^43, 44^, we hypothesized that the P21C9 mutant might be sensitive to AMPs due to reduced PGN O-acetylation as a consequence of reduced secretion of acyltransferases. Consistent with our hypothesis, we detected significantly reduced PGN acetylation levels in the P21C9 mutant. An *L. plantarum* mutant lacking both acyltransferases^43^ showed the expected lack of PGN acetylation, validating our assay (Fig. 4h). Importantly, we could rescue PGN acetylation in the P21C9 mutant to wild-type level by *ffh* and *oatA* overexpression (Fig. 4i). *OatA* overexpression also reduced the binding of cationic AMPs to P21C9 cells (Fig. 4j) and rescued P21C9 mutant sensitivity to polymyxin B (Fig. 4k). This suggests, that the absence of ffh leads to a decreased secretion of the oatA protein, which is responsible for the low PGN acetylation, negative surface charge, and increased sensitivity to AMPs in the P21C9 mutant. Consistent with this, *L. plantarum* mutants lacking acyltransferases showed increased binding to 5-FAM-LC-LL37 (Fig. 4l) and enhanced susceptibility to polymyxin B (Fig. 4m). Together these results establish O-acetylation as a new PGN modification mediating bacterial sensitivity to cationic AMPs via surface charge alternations.

**Figure 4.**
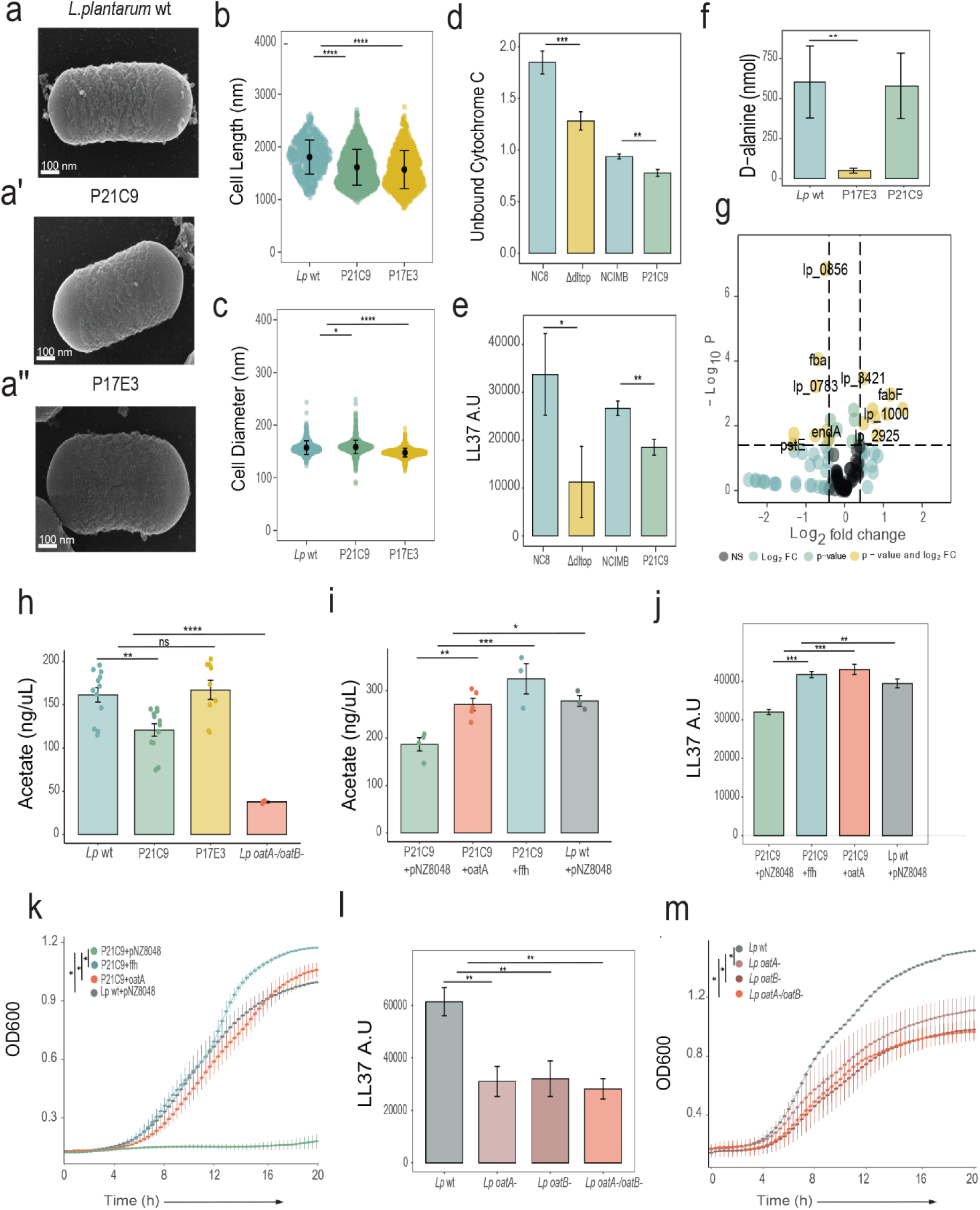
Microbiota resistance to the AMPs is driven by O-acetylation of peptidoglycan and D-alanylation of the teichoic acid. **a**, Scanning electron microscopy image of *L. plantarum^NCIMB^*, P21C9 (**a’**), and P17E3 (**a”**) cell. **b**, Cell Length and **c**, Cell Diameter of *L. plantarum^NCIMB^* (n=612), P21C9 (n=1535), and P17E3 (n=1006). Individual dots show single cell record. Violin dot plots show median and interquartile ranges. **d-e**, Binding of *L. plantarum^NCIMB^*, P21C9, and P17E3 to Cytochrome C (**d**) and to labelled antimicrobial peptide LL37 (**e**) (n= 3 independent experiments). **f**, HPLC quantification of D-alanine released by whole cells of *L. plantarum^NCIMB^*, P21C9, and P17E3 (n=6 independent cultures). Bar plots show Mean and SEM. **g**, Differential analysis of secreted and membrane-bound proteins in *L. plantarum^NCIMB^* and P21C9. Lines in volcano plot show the p-value cut-off (0.05) and 0.5 as FC (n=3 independent samples). **h**, Quantification of acetate released from peptidoglycan extracted from *L. plantarum^NCIMB^* (n=12), P21C9 (n=12), P17E3 (n=9), and *L. plantarum^oatA-/oatB-^* (n=3). **i**, Quantification of acetate released from peptidoglycan extracted from P21C9 mutant overexpressing *oatA* (n=5) or *ffh* (n=3). P21C9 mutant (n=4) and wild type *L. plantarum* (n=3) containing empty pNZ8048 plasmid were used as controls. Bar plots show Mean and SEM. **j**, Binding of the indicated strains to labelled antimicrobial peptide LL37 (n= 3 independent experiments). **k**, Kinetics of the growth of P21C9+pNZ8048 (empty plasmid), P21C9+*ffh*, P21C9+*oatA*, and *L.plantarum^NCIMB^* + pNZ8048 in MRS media supplemented with Polymyxin B (n= 3 independent experiments). Mean and SEM are shown. **l**, Binding of *L. plantarum^NCIMB^*, *L. plantarum^oatA-^*, *L. plantarum^oatB-^*, and *L. plantarum^oatA-/oatB-^*to labelled antimicrobial peptide LL37. Bar plots show Mean and SEM (n= 3 independent experiments). **m**, Kinetics of the growth of *L. plantarum^NCIMB^*, *L. plantarum^oatA-^*, *L. plantarum^oatB-^*, and *L. plantarum^oatA-/oatB-^*in MRS media supplemented with Polymyxin B, (n= 3 independent experiments). Mean and SEM are shown. *P < 0.05, **P < 0.01, ***P < 0.001, ****P < 0.0001. Kruskal–Wallis and Bonferroni post hoc tests were used for statistical analysis. Simple linear regression analysis was performed for the kinetics analysis.

### Resistance to AMPs is essential for *L. plantarum* persistence in the gut during immune activation

Next, we investigated the persistence of AMP-sensitive *L. plantarum* mutants in the *Drosophila* gut. First, we used gnotobiotic flies monocolonized with wild-type *L. plantarum* and *Δdltop* mutant and measured their abundance in uninfected and *Ecc15*-infected flies (Fig. 5a). The load of the *Δdltop* mutant was significantly lower compared to wild-type *L. plantarum* 6h and 24h post *Ecc15* infection. However, the *Δdltop* mutant colonized uninfected flies as efficiently as wild-type *L. plantarum*, suggesting that the decline after infection is not due to a general incapacity to persist in the gut, but rather to sensitivity to immune activation. Supporting this, the load of the *Δdltop* mutant did not decline after infection in flies lacking AMPs (Δ*AMP)*. Second, using a priming approach (Fig. 5b), we confirmed that the reduction of the *Δdltop* mutant abundance after infection is rescued in Δ*AMP* flies. Finally, genetic overactivation of the Imd pathway in the gut resulted in a significant decrease in *Δdltop* mutant levels compared to control flies and wild-type *L. plantarum* at all time points tested (Fig. 5c). The P21C9 mutant behaved the same – it stably colonized uninfected flies but failed to persist in the gut after *Ecc15* infection or genetic activation of the immune response (Supplementary Fig. 3a-c). Persistence of the P21C9 mutant during infection was restored in Δ*AMP* flies, proving that AMPs at least in part are responsible for clearance of AMP-sensitive mutants during infection. Overall, these results indicate that resistance to host AMPs is essential for *L. plantarum* to stably persist in the inflamed gut environment.

**Figure 5.**
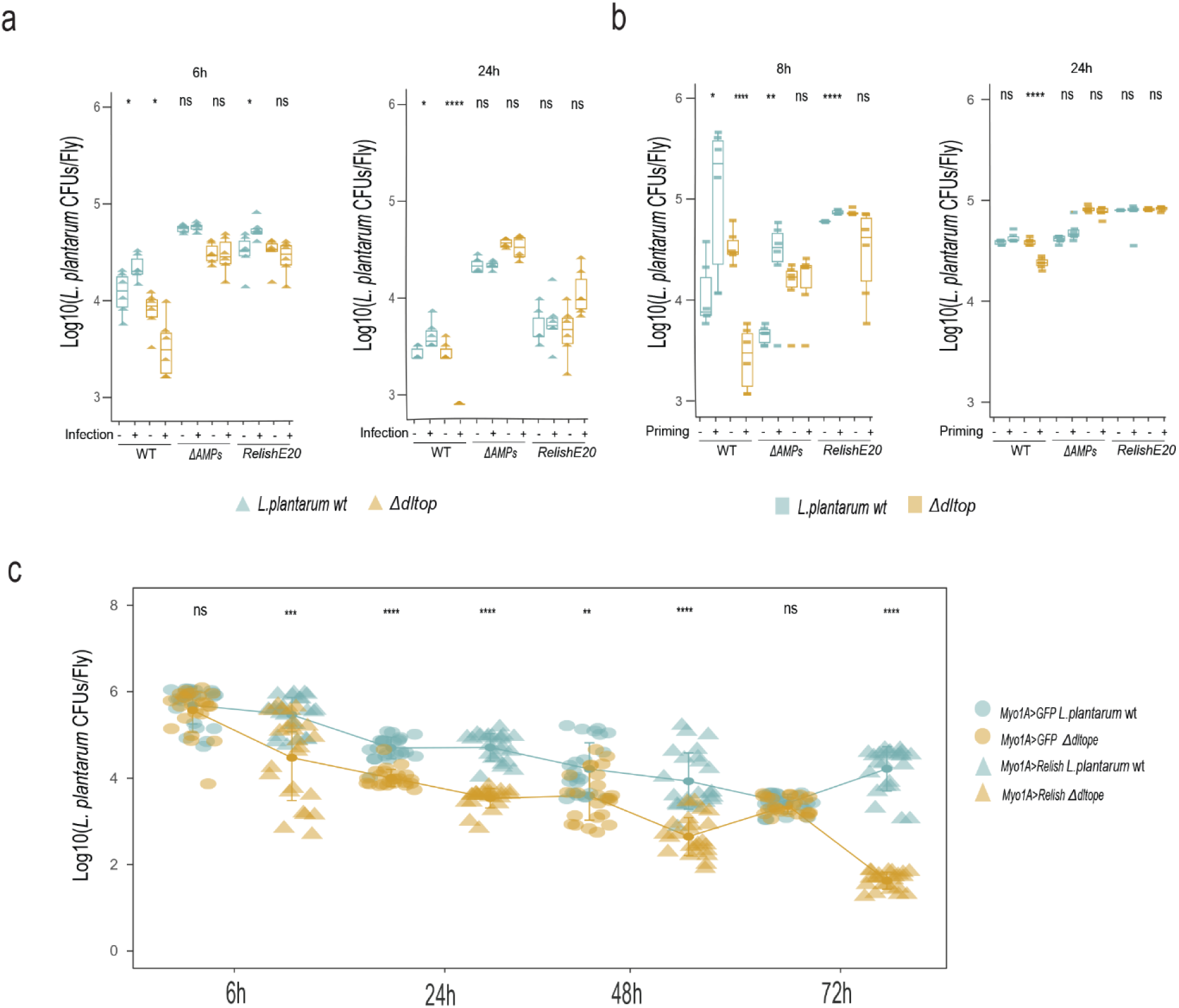
*L. plantarum delta* operon genes are essential to colonize and persist in the gut during infection. **a**, *L. plantarum^NC8^* and *L. plantarum^Δdltop^* loads in wild type, Δ*AMPs* and *Relish^E20^* flies after the infection with *Ecc15* at 6h and 24h (n=6 independent samples per treatment with 5 flies per sample). **b**, *L. plantarum^NC8^* and *L. plantarum^Δdltop^* loads after the priming with *Ecc15* at 8h and 24h in wild type, Δ*AMPs* and *Relish^E20^* (n=6 independent samples per treatment with 5 flies per sample). **c**, *L. plantarum^NC8^* and *L. plantarum^Δdltop^* loads in *Myo1A-GAL4>UAS-GFP* and *Myo-GAL4>UAS-Relish* flies at 6h, 24h, 48h, and 72h after colonization (n=24 samples per treatment with 5 flies per sample). Individual tringles, squares and dots show mean CFU values from pools of n = 5 animals in the Log10 scale. Boxplots and dot plots show median and interquartile ranges (IQR); whiskers show either lower or upper quartiles or ranges. *P < 0.05, **P < 0.01, ***P < 0.001, ****P < 0.0001. Kruskal–Wallis and Bonferroni post hoc tests were used for statistical analysis.

### Resistance to AMPs is essential for *L. plantarum* persistence in the gut during aging

Finally, we investigated what happens with long-term persistence of AMP-sensitive mutants and how they impact aging phenotypes. The main motivation to investigate such age-related phenotypes is that the microbiota load is known to increase with age despite increased expression of AMPs. We hypothesized that microbiota resistance to host AMPs is essential to survive this age-associated immune activation. To test this hypothesis, we colonized wild-type, *relish*, Δ*AMP* flies with wild-type and *Δdltop* bacteria, respectively and scored their lifespan, bacterial load, and several aging hallmarks. Wild-type flies colonized with the *Δdltop* mutant lived longer compared to flies colonized with wild-type bacteria. Both Δ*AMP* and *relish* mutants were short-lived compared to wild-type flies and there was no significant difference in the lifespan between wild-type and *Δdltop* mutant-colonized treatments (Fig. 6a). As expected, the bacterial load increased with age in all tested treatments. However, while the *Δdltop* mutant reached significantly lower density in the guts of wild-type flies, it colonized guts of Δ*AMP* and *relish* mutants as efficiently as the wild-type strain (Fig. 6b), suggesting that AMPs reduce *Δdltop* load in wild-type flies. Consistent with the observed lifespan and bacterial load, Δ*AMP* and *relish* mutants exhibited excessive stem cell proliferation (as measured by PH3 staining) particularly after colonization with the *Δdltop* mutant (Fig. 6c). Such intestinal dysplasia was significantly reduced in *Δdltop-*colonized wild-type flies. The JAK-STAT pathway, as one of the major drivers of stem cell proliferation, showed increased activation with aging in the guts of *L. plantarum* wild-type-colonized flies. Similar results were also observed for the activation of the Imd pathway (Fig. 6d). These results suggest that AMPs are responsible for controlling commensal load during aging, consistent with the findings of Hanson et al,^45^ and that resistance to AMPs is essential for commensals persistence in the inflamed gut environment of aging flies.

**Figure 6.**
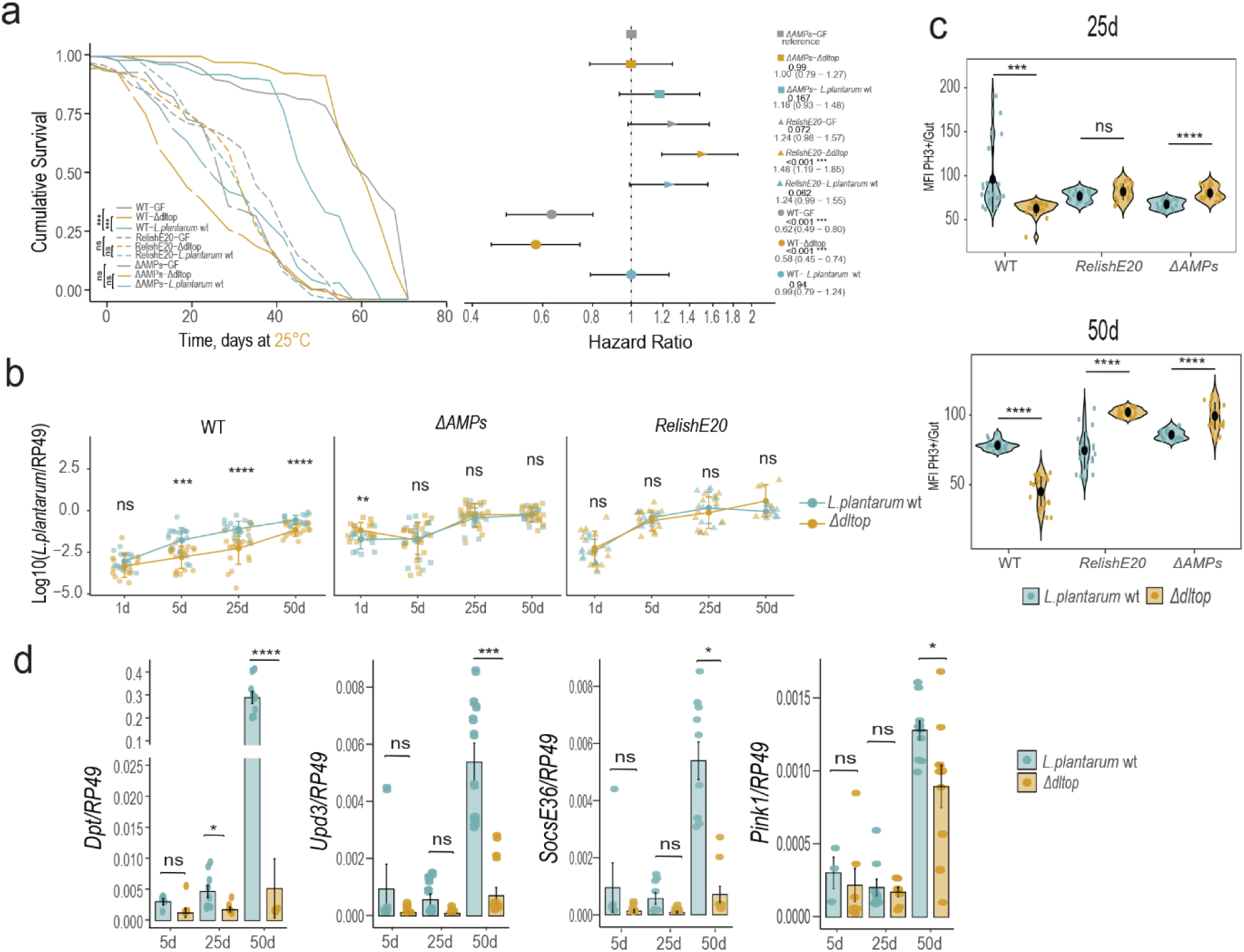
*L. plantarum* resistance to the AMPs is a crucial regulator of the onset of *Drosophila* aging. **a**, on the left panel Kaplan–Meier survival curves of mono-colonized flies with either *L. plantarum^NC8^* or *L. plantarum^Δdltop^*, germ-free flies were used as control. P values were obtained from stratified groups of fly genotypes, the log-rank test was applied to the survival curves per stratified group (n=3). **a**, right panel shows Hazard ratios based on a Cox proportional-hazards model in mono-colonized flies. Forest plot shows 3 independent experiments, horizontal bars show 95% confidence intervals (CI), P values were obtained from z tests for individual sample. **b**, *L. plantarum^NC8^*(n1) or *L. plantarum^Δdltop^* (n2) loads in wild type 1d- (n1=15, n2=16), 5d- (n1=20, n2=19), 25d- (n1=19, n2=19), 50d-old (n1=16, n2=16); in *ΔAMPs* 1d- (n1=14, n2=14), 5d- (n1=20, n2=19), 25d- (n1=18, n2=20), 50d-old (n1=14, n2=18), and in *Relish^E20^* 1d- (n1=13, n2=13), 5d- (n1=11, n2=14), 25d- (n1=12, n2=11), 50d-old (n1=9, n2=7) mono-colonized flies. Individual dots show mean load values determined by qPCR from pools of n = 5 animals in the Log10 scale. **c**, Intestinal Stem Cell proliferation, indicated by Median Fluorescence Intensity (MFI) of Phospho-histone H3-positive cells in wild type, *ΔAMPs,* and *Relish^E20^* mono-colonized flies with *L. plantarum^NC8^* or *L. plantarum^Δdltop^* at 25d or 50d (n=21 guts per sample). **e**, Gene expression of *DptA*, *Upd3*, *Socs36E*, and *Pink1* in 5d- (n1=5, n2=7), 25d- (n1=9, n2=7), and 50d- (n1=10, n2=10) wild type flies mono-colonized with *L. plantarum^NC8^*or *L. plantarum^Δdltop^*. Individual dots show gene expression per 20 guts of female flies. Bar plots show Mean and SEM. Violin plots show median and interquartile ranges (IQR). *P < 0.05, **P < 0.01, ***P < 0.001, ****P < 0.0001. Kruskal–Wallis and Bonferroni post hoc tests were used for statistical analysis.

## Discussion

In this study, we show that infection – one of the frequent perturbations occurring in the digestive tract, had little impact on *Drosophila* microbiota composition and abundance. We identified resistance to AMPs as a key feature of microbiota resilience during intestinal inflammation. Thus, our work reveals that in addition to host immune tolerance to the microbiota, commensal-encoded resilience mechanisms are necessary to maintain a stable host-microbiota association during inflammation.

The intestinal immune response is regionalized in *Drosophila* and other insects^31^. This regionalization leads to the formation of gut regions with strong expression of negative regulators of the immune response, thus creating a protective zone for symbiotic microbes^19, 46^. Moreover, many insects are equipped with specialized symbiotic organs – bacteriomes, where symbionts are maintained. Bacteriomes allow the host to create a favourable environment for symbionts but also to keep them under control and protect them from perturbations, like infections^47^. While compartmentalization is an efficient strategy to protect the symbionts from the immune response, we could detect *L. plantarum* in gut regions with strong AMP expression, suggesting that *L. plantarum* does not hide from the effectors in protective gut zones. Instead, our results support a hypothesis that resistance to AMPs mediates commensal resilience during gut inflammation. Using a genetic screen in *L. plantarum*, we identified several determinants of AMP resistance. One of the identified AMP-sensitive mutants had a transposon insertion in the *dlt* operon, which has been previously implicated in the sensitivity to AMPs in several Gram-positive bacteria^37^. Consequently, *dlt* mutants in several pathogens exhibit attenuated virulence^23, 48, 49^. The *dlt* operon was also found to be essential for the commensal establishment in the host gut, namely for *L. reuteri* in the mouse intestine^50^ and for *L. casei* in the rabbit gut^51^. Attieh et al reported that the *L. plantarum dlt* mutant is impaired in the colonization of the *Drosophila* gut and that it triggers a stronger immune response compared to wild-type *L. plantarum*^23^. Our results similarly support the essential role of the *dlt* operon in *L. plantarum* in colonizing the *Drosophila* gut. However, it was especially important during an immune challenge because the *dlt* operon mutant colonized uninfected flies as efficiently as wild-type *L. plantarum*. A functional *dlt* operon is also required for *L. plantarum* to promote larval growth of *Drosophila* under chronic undernutrition^39^, illustrating multiple essential roles of WTA D-alanylation in host-microbe interactions. It is intriguing that the same cell wall modification is crucial for the stable persistence of bacteria in the gut, immunomodulation, and to confer a beneficial impact on the host. This pleiotropy of WTA D-alanylation results in the scenario where commensals lacking D-alanylated WTAs are better sensed by PGRPs and trigger a strong immune response which will eventually eliminate such commensals due to their sensitivity to AMPs. Commensals with D-alanylated WTAs, however, will promote host tolerance mechanisms and will establish a stable association with the host. Thus, D-alanylated WTAs might be used as a signal to recognize beneficial commensals and trigger either tolerance or immune response.

In addition to the *dlt* operon, we identified a new mediator of bacterial sensitivity to AMPs – *ffh,* which is an integral part of the signal recognition particle pathway (SPR). Apart from being important for virulence^52^, the role of the SPR pathway in microbial interactions with the host has not been studied. Such a role is very likely, considering that SPR pathway mutants in different bacteria are impaired in the secretion and translocation of proteins necessary for adhesion, biofilm formation, PGN and cell wall biosynthesis^40–42^. Disruption of the SPR pathway likely has a pleiotropic effect on bacterial physiology and could alter sensitivity to AMPs in several ways. Our results, however, support a prominent role of reduced PGN O-acetylation due to reduced secretion of acyltransferases in the sensitivity of the P21C9 (*ffh*) mutant to AMPs. While PGN O-acetylation is a well-established mechanism of bacterial resistance to lysozyme^44^, we found that it also mediates sensitivity to cationic AMPs likely by altering the surface charge. It would be intriguing to explore the extent to which the role of PGN O-acetylation in AMP sensitivity is conserved among bacteria.

Earlier studies showed an essential role of AMPs in shaping *Drosophila* gut communities. For instance, Marra et al found that the abundance of multiple commensals, particularly *Acetobacter sp*, is increased in Δ*AMP* mutant flies, supporting a prominent role of AMPs in controlling *Acetobacter* species^28^, which, as we showed here, are more sensitive to AMPs than other fly commensals. Despite the increased sensitivity, *A. melorum* still stably colonized flies during infection, suggesting that commensals rely on additional mechanisms besides resistance to AMPs to survive in an inflamed host environment. These mechanisms, which could include priming by AMPs^53^, remain to be investigated. Previously, the role of AMPs in controlling the abundance of *L. plantarum* in the fly gut was found to be less pronounced and was evident only in *L. plantarum* monoassociated flies but not when additional community members were present^28^. We also observed elevated *L. plantarum* abundance in some cases in Δ*AMP* mutants, suggesting that while *L. plantarum* is resistant to some cationic AMPs like polymyxin B and CecB, it is very likely that there are AMPs or their combinations to which *L. plantarum* is sensitive. Overall, our results support the current view that AMPs control gut commensals. However, some of the microbiota members are more resistant to AMPs, allowing these microbes to better colonize the host during infection. In turn, the host likely relies on additional means to control these commensals and maintain a balanced microbiome. Lysozymes and ROS are among the likely suspects that would be interesting to investigate in the future^54^.

Collectively, our work shows that AMP resistance via cell wall modifications historically associated with pathogen virulence is also essential to maintain stable microbiota-host interactions. This adds more evidence that host-symbiont and host-pathogen associations are mediated by the same molecular dialog^55^. Further elucidation of the mechanisms of microbiota resilience during inflammation and generality of such mechanisms will be an exciting future endeavour.

## Materials and Methods

### *Drosophila* stocks and rearing

The following *Drosophila* stocks used in this study were kindly provided by Dr. Bruno Lemaitre: DrosDel *w^1118^* iso; *Relish^E20^* iso; *ΔAMP* iso; *UAS-Relish*; *UAS-CecA*; *w;Myo1A-Gal4,tub-Gal80TS,UAS-GFP*; *w;esg-Gal4,tub-Gal80TS,UAS-GFP*^13,22, 45^. The following stocks were obtained from the Bloomington Drosophila Stock Center: *w*; P{DptA-GFP.JM863}D3-2 P{DptA-GFP.JM863}3-4* (55709); *UAS-mCD8::GFP* (32185). The stocks were routinely maintained at 25 °C with 12/12 h dark/light cycles on a standard cornmeal-agar medium: 3.72g agar, 35.28g cornmeal, 35.28g inactivated dried yeast, 16 ml of a 10% solution of methyl-para-ben in 85% ethanol, 36 ml fruit juice, 2.9 ml 99% propionic acid for 600 ml. Food for germ-free flies was supplemented with Ampicillin (50 µg/ml), Kanamycin (50 µg /ml), Tetracyclin (10 µg/ml), and Erythromycin (10 µg /ml). Fresh food was prepared weekly to avoid desiccation. Female flies were used for all experiments.

### Generation of germ-free flies

To obtain axenic fly stocks, embryos laid over a 16-h period on grape juice plates were collected from 4-to 10-day-old females. Embryos were rinsed in phosphate-buffered saline (PBS) and transferred to 1.5 ml tubes. All following steps were performed in a sterile hood. Embryos were placed in a 3% solution of sodium hypochlorite for 10 min. The bleach solution was discarded and embryos were rinsed three times in sterile PBS. Embryos were transferred by pipette to tubes with antibiotics-supplemented food in a small amount of 100% ethanol and maintained at 25 °C. Subsequent generations were maintained in parallel to their conventionally reared counterparts by transferring adults to new tubes with antibiotics-supplemented food. The axenic state of flies was routinely assessed by culturing.

### Bacterial strains and culture conditions

The strains used in this study are listed in Supplementary Table 2. *Escherichia coli* strains and *P. aeruginosa* were grown at 37 °C in LB medium with agitation. *L. plantarum* strains were grown in static conditions in MRS medium at 37 °C, unless stated otherwise. *Ecc15* and *P. entomophila* were grown at 30 °C in LB medium with agitation. *E. faecalis* was grown at 37 °C in BHI medium with agitation. *A. pomorum* was grown at 30 °C in MRS medium with agitation. *A. malorum, Sphingomonas sp, Variovorax sp, G. morbifer* were grown at 30 °C in mannitol medium with agitation. Erythromycin antibiotic was used at 5 μg/ml for *L. plantarum* and 150 μg/ml for *E. coli*. Chloramphenicol was used at 10 μg/ml. Solidified media were used for estimation of colony forming units (CFUs).

### Generation and screening of random transposon mutant library in *L. plantarum* NCIMB8826

*L. plantarum* transposon mutagenesis was performed using the P_junc_-TpaseIS*_1223_* system as previously described^34, 35^. Briefly, electrocompetent *L. plantarum* NCIMB8826 cells were first transformed with pVI129, resulting in the NCIMBpVI129 strain. Electrocompetent cells of the NCIMBpVI129 strain were transformed with pVI110, plated on MRS plates supplemented with 5 μg/ml of erythromycin and incubated for 48 h at 42 °C to select for integrants. A total of 3000 mutants were individually stored at −80 °C in deep 96-well plates. The whole transposon library was screened in 96-well plates for mutants not able to grow in MRS supplemented with 50 µg/ml of polymyxin B.

### Mapping of transposon insertion sites

Genomic DNA from transposon mutants was digested sequentially with ClaI and BstBI restriction enzymes (NEB). These enzymes create compatible sticky ends facilitating ligation. Digested fragments were ligated using T4 DNA ligase (Thermo Fisher) according to the manufacturer’s instructions. The resulting ligation products were transformed into the *E. coli* TG1 strain, in which circularized fragments that contain the transposon replicate as plasmids. Plasmids were isolated from selected transformants using Monarch plasmid miniprep kit (NEB) and subjected to sequencing reactions (Eurofins) using the primers IRR6 and IRL6, which target the transposon sequence extremities. Identification of transposon target sequences was performed with the BLAST software from the National Center for Biotechnology Information (http://blast.ncbi.nlm.nih.gov/). Primers used in this study are listed in Supplementary Table 3.

### DNA extraction and 16S rRNA analysis

10d old *w^1118^* iso conventional flies were infected with alive or heat-killed *Ecc15*, OD=200 mixed 1:1 with 5% sucrose. 150 µl of the mixture was applied to a filter disk covering the food surface of the vial with food; control flies were supplemented with 2.5% sucrose. Flies were incubated at 29 °C for 6h and 24h. 20 guts per sample were collected, and the total DNA was extracted using Puregene kit (Qiagen). Briefly, the guts were smashed in 300 µl of cell lysis solution with a pestle, in 1.5 ml Eppendorf tubes, and incubated 15 min at 65 °C. 100 µl of protein precipitation solution was added to each sample, vortexed, and incubated for 5 min on ice. Samples were spun down for 5 min at max speed; supernatants were transferred into new 1.5 ml Eppendorf tubes with 300 µl of cold isopropanol and pelleted at max speed for 10 min. Pellet was washed with 70% ethanol and spun down for 1 min at max speed. The dried pellet was dissolved in 20 µl of DNA hydration buffer. V3-V4 region of 16S rRNA gene was amplified using specific primers with the barcode and Phusion High-Fidelity PCR Master Mix (New England Biolabs). PCR products were mixed at equal density ratios. The mixed PCR products were purified with Qiagen Gel Extraction Kit (Qiagen, Germany). The libraries were generated with NEBNext UltraTM DNA Library Prep Kit for Illumina and quantified via Qubit and qPCR. The libraries were sequenced on NovaSeq 6000 system in paired end mode (Novogene). Analysis was performed using QIIME^56^.

### *Drosophila* mono-colonization-infection, priming-colonization, mono-colonization and infection in overexpression

**(a)** mono-colonization-infection: 10d old germ-free flies were starved for 2h prior to mono-colonization with *L. plantarum* or *A. malorum*. Overnight bacterial cultures were adjusted to OD=50 and mixed 1:1 with 5% sucrose. 150 µl of the mixture was placed on paper filter disks covering fly food and incubated at 25°C. After 48h of colonization, flies were starved for 2h at 29°C. After starvation, flies were infected with 150 µl of either alive or heat-killed *Ecc15*, OD=200 mixed 1:1 with 5% sucrose; control flies were treated only with 2.5% sucrose. Infection was performed at 29°C. 5 flies per treatment were used to record bacteria loads in MRS, Mannitol, or by qPCR, using total DNA at 6h and 24h post-infection. **(b)** Priming-colonization: 10d old germ-free flies were starved for 2h and primed with 150 µl of either alive or heat-killed *Ecc15* at 29°C; control flies were treated only with sucrose 2.5%. After 3h of priming, flies were starved for 2h and mono-colonized with *L. plantarum* and *A. malorum* or infected with *Ecc15* or *Pe*, OD=200 mixed 1:1 with 5% sucrose. Control flies were treated only with 2.5% sucrose. Bacteria loads were recorded at 8h and 24h after priming using 5 flies per treatment. Bacteria loads were determined in MRS or Mannitol for microbiota and LB for pathogens or by qPCR, using total DNA. **(c)** mono-colonization and infection in overexpression: Before colonization or infection, flies were maintained and starved for 2h at 29°C. 150 µl of *L. plantarum* or *A. malorum* OD=50 mixed 1:1 with 5% sucrose was added to a paper filter disk. For Infection, 150 µl of *Ecc15* OD=200 mixed 1:1 with 5% sucrose was used. Bacteria loads were recorded by CFUs or qPCR at 6h, 24h, 48h, and 72h. Flies were flipped into conventional vials 48h post-treatment.

### Aging and Survival

3d-old female flies were mono-colonized either with *L. plantarum* NC8 or *L. plantarum* Δ*dltop.* Bacteria loads were recorded by qPCR. 20 guts per sample were dissected to record the gene expression. Survival was recorded every 2 days and flies were flipped into new conventional vials every counting day.

### Imaging and Immunostaining

DptA-GFP germ-free flies were mono-colonized with *L. plantarum* WJL-mCherry for 24h and infected with *Ecc15*. After 6h guts were fixed in PBS 1X with 4% paraformaldehyde. DNA was stained with DAPI 5 mg/ml (Sigma-Aldrich) 1:1000 in 1ml PBS overnight at 4°C, rinsed once with PBS 1X, and mounted in a chamber with Mowiol 4% (Sigma-Aldrich). Images were obtained using PS8 lightning confocal microscope. PH3 staining: 25d and 50d old guts were incubated in PBT 0.1% Triton X-100 + 0.3% BSA and 1:500 anti-PH3 mouse mAb IgG1 overnight at 4°C, rinsed in PBT and stained with secondary antibody Alexa555-coupled goat anti-rabbit antibody 1:500 and DAPI 5 mg/ml, 1:1000 at RT 1h in the Belydancer, rinsed in PBS and mounted in a chamber with Mowiol 4%. The gut of Esg-GAL4, UAS-GFP flies were dissected in PBT 0.1% Triton X-100 + 0.3% BSA and stained in DAPI 5mg/ml, 1:1000, after incubation of 1h RT, the guts were mounted in with Mowiol 4%. Images were obtained using a Leica DMR Fluorescence microscope.

### Regionalization of *Drosophila* gut

*w^1118^* iso germ-free flies were mono-colonized with *L. plantarum* for 24h and infected with *Ecc15* for 6h. 80 guts per sample were dissected on ice and cut into regions R0 to R5. Regionalization of the gut was determined according to Buchon et al., 2013^57^. Samples of gut regions were smashed in 100 µl of PBS 1X and split into 50 µl for DNA extraction and RNA extraction. Gene expression and bacteria loads were determined by qPCR in paired samples.

### Minimal Inhibitory Concentration (MIC) and growth kinetics

Overnight bacterial cultures were adjusted to OD 0.1 and diluted 1:100 and cultured in 96 well plates with Polymyxin B or Cecropin B. Next day, the bacterial growth was recorded as MIC value. The overnight culture was adjusted to 0.05 and grew 20h in 96 well plates in a plate reader at 37°C, in MRS medium supplemented with Polymyxin B. OD600 was measured to determine the kinetics of Bacteria in presence of AMP *in vitro*. Reads were performed in infinite 200 Pro plate reader (Tecan).

### Cytochrome C and 5-FAM-LC-LL37 binding

Bacterial cultures were grown to OD 0.6 and resuspended in Buffer A (KH2PO4 1M pH: 7.0, BSA 0.01%) and Cytochrome C (Sigma) solution (0.5 mg/ml), the cells were incubated 15 min RT. Samples were centrifugated, and supernatants were measured in 96 well plates at 440 nm. Overnight cultures were adjusted to OD 0.1 in PBS 1X, and 5-FAM-LC-LL37 (Eurogentec) solution (14 µM) was added to each sample and incubated for 1h at 37°C, 590rpm. Samples were centrifugated for 1 min at max speed and supernatants were transferred to 96 well plates. Fluorescence was measured at absorbance 494 nm and 521 nm of emission. Reads were performed in infinite 200 Pro plate reader (Tecan).

### RNA extraction and RT-qPCR

RNA was extracted from 20 guts per treatment using TRIzol reagent as previously described^58^. RNA concentration was determined by Nanodrop ND-1000 spectrophotometer. RT-PCR was performed using 500 ng of RNA in 10 µl volume of Solution with PrimeScript RT (TAKARA) and random hexamer primers. Quantitative PCR was performed in 384-well plates using the SYBR Select Master Mix from Applied Biosystems. Reads were performed on a LightCycler 480 (Roche).

### Samples preparation for proteomics

Bacteria were cultured to OD 0.5. Cells were removed by centrifugation for 15 min at 3600 rpm. Supernatants were filtered through 0.22 µm filter to remove any remaining bacteria. 20 ml of supernatants were placed in Macrosep Advance Centrifugal Devices MWCO 3 kD (Pall Corporation) and centrifuged at max speed at 4°C for 3h. The concentrated supernatant solutions were transferred into Eppendorf tubes and sent to High Throughput Mass Spectrometry Core Facility (Charité) for proteomic analysis. The differential analysis of quantitative proteomics data was performed in Perseus v2.0.7.0.

### Isolation and measurement of peptidoglycan acetylation

*L. plantarum* PGN was extracted as described before^59^. Briefly, *L. plantarum* cells were grown in MRS medium and harvested by centrifugation at mid-exponential phase. Collected cells were adjusted to OD 10 and 2 ml of this cell suspension were transferred to 2 ml centrifuge tube. The cells were collected by centrifugation, washed with MilliQ water, boiled for 10 min and collected by centrifugation again. The pellet was resuspended in 1 ml of 5% SDS in 50 mM of MES (Sigma-M8250) pH 5.5, and boiled for 25 min at 100°C in a heating block. The pellets were collected by centrifugation, resuspended in 1 ml of 5% SDS in 50 mM of MES (Sigma-M8250) pH 5.5, and boiled for 15 min and then washed twice with MilliQ water to remove SDS traces. Next, the pellets were sequentially enzymatically treated with 2 mg/ml of Pronase (Roche 165921) in 50 mM of MES (Sigma-M8250) at pH 6.0 for 90 min at 60 °C; 200 µg/ml of trypsin (Sigma T-0303) in 50 mM of MES (Sigma-M8250) at pH 6.0 for 2 h at 37 °C with shaking; DNase (Sigma-D-4527) and RNase (Sigma-R-5503) (50 µg/ml) in 50 mM of MES (Sigma-M8250) at pH 6.0 for 1 h at 37 °C. After centrifugation, the pellets were resuspended in 2% SDS in 50 mM of MES (Sigma-M8250) at pH 5.5 and boiled for 15 min. Finally, the pellets were washed with MilliQ water to remove SDS traces, lyophilized and stored at −20 °C. Measurement of Peptidoglycan O-Acetylation was performed as described previously^60^ with certain modifications. Briefly, lyophilized cell wall extracts (the same amount by weight per sample) were resuspended with 500 µl of 500 mM NaOH and incubated at room temperature for 30 minutes to release any ester-linked acetate. Solutions were neutralized with 500 mM H_2_SO_4_ and subjected to centrifugation. Quantification of released acetate was performed using Acetate Colorimetric Assay Kit (MAK086, Sigma), following manufacturer’s instructions.

### Quantification of D-alanylation of teichoic acids

D-alanylation of teichoic acids was quantified as described previously^39, 61^. Briefly, bacterial overnight cultures were centrifuged and resuspended in 1 ml of 1X PBS, washed with 1 ml of acetate ammonium (20 mM pH: 4.7). Solution was split into 1.5 ml Eppendorf tubes at OD 50 and heat-inactivated at 100 °C for 10 min. Cells were frozen with nitrogen stream for 1 min and lyophilized in the vacuum chamber overnight. d-alanine was released from heat-inactivated bacteria by mild alkaline hydrolysis with 0.1 N NaOH for 1 h at 37 °C. Finally, the solutions were neutralized with 0.1 N HCl. After neutralization, the extracts were incubated with Marfey’s reagent (1-fluoro-2,4-dinitrophenyl-5-L-alanine amide; Sigma). After drying in a nitrogen stream, the residues were derivatized with 50 µL of 0.5% Marfey’s reagent (w/v in acetone) and 100 µL of 125 mM disodiumtetraborate for 30 min at 40 °C. The reaction was stopped with 25 µL pf 4 M HCl. The resulting solution was diluted (1:10) with 4 mM ammonium formiate, pH 4.6). Two µL of the derivatized samples were subjected on an Acquity H-Class UPLC system (Waters), using an AccQ-Tag Ultra C18 column (1.7µm, 2.1×100mm), and a linear gradient of 100% mobile phase A (4mM ammonium formiate; pH 4.6) to 50% mobile phase A and B (acetonitrile) within 5 min at 0.5 mL/min. Derivatized amino acids were detected both, by UV-absorbance (340 nm) and by mass spectrometry (negative ion mode at m/z 340) with a single quadrupole mass spectrometer (QDA, Waters). For quantification, standard D-alanine was analysed at 5 concentrations (0-100µM) which were prepared in duplicate in 0.1 N HCl before derivatization.

### Scanning electron microscopy

Bacterial cells were fixed with 2.5% glutaraldehyde and 20 µl drops of bacterial suspension were spotted onto polylysine-coated round glass coverslips placed into the cavities of a 24-well cell culture plate. After 1 h of incubation in a moist chamber, PBS was added to the wells, and the samples were fixed with 2.5% glutaraldehyde for 30 min. Samples were washed and post-fixed using repeated incubations with 1% osmium tetroxide and 1% tannic acid, dehydrated with a graded ethanol series, critical point dried and coated with 3 nm platinum/carbon. Specimens were analyzed in a Leo 1550 field emission scanning electron microscope using the in-lens detector at 20 kV. For quantification, images were recorded at a magnification of 2000 x and analyzed with the Volocity 6.5.1 software package.

### Statistical analysis

Statistical parameters and tests are shown in the respective figure legends. Boxplots display boxes with the interquartile range from first to third quartiles; whiskers show the tenth and ninetieth percentiles. Statistical test was performed using R v4.2.2. Survival analysis and individual median survival were performed using Kaplan-Meier method and Log Rank test survival with the R package survminer. Comparisons were performed in pairs and plotted together. The R packages ggplot2, dplyr, and tidyverse were used for data visualization.

## Supporting information

Combined supplementary information

## Acknowledgements

We are grateful to Bruno Lemaitre and the Bloomington Drosophila Stock Center (NIH P40OD018537) for fly stocks. We thank Pascale SERROR (Michalis Institute) for sharing pVI110 and pVI129 plasmids, Pascal Hols (Université catholique de Louvain) for sharing pNZ8048 plasmid, Marie-Pierre CHAPOT-CHARTIER (Michalis Institute) for kindly providing *L. plantarum* mutants lacking PGN acetyltransferases, François Leulier and Renata Matos for sharing *L. plantarum* WJL-mCherry, *L. plantarum* NC8 and *Δdltop* mutant. We thank the Core Facility High Throughput Mass Spectrometry of the Charité for support in acquisition and analysis of the proteomics data. We thank Alexandra Hrdina for editing of the manuscript. This work was supported by the Max Planck Society.

## Notes

### Competing Interest Statement

The authors have declared no competing interest.

