## Supplementary material for "Resistance to host antimicrobial peptides mediates resilience of gut commensals during infection and aging in *Drosophila*": Combined supplementary information

To manuscript

**Supplementary figures:**

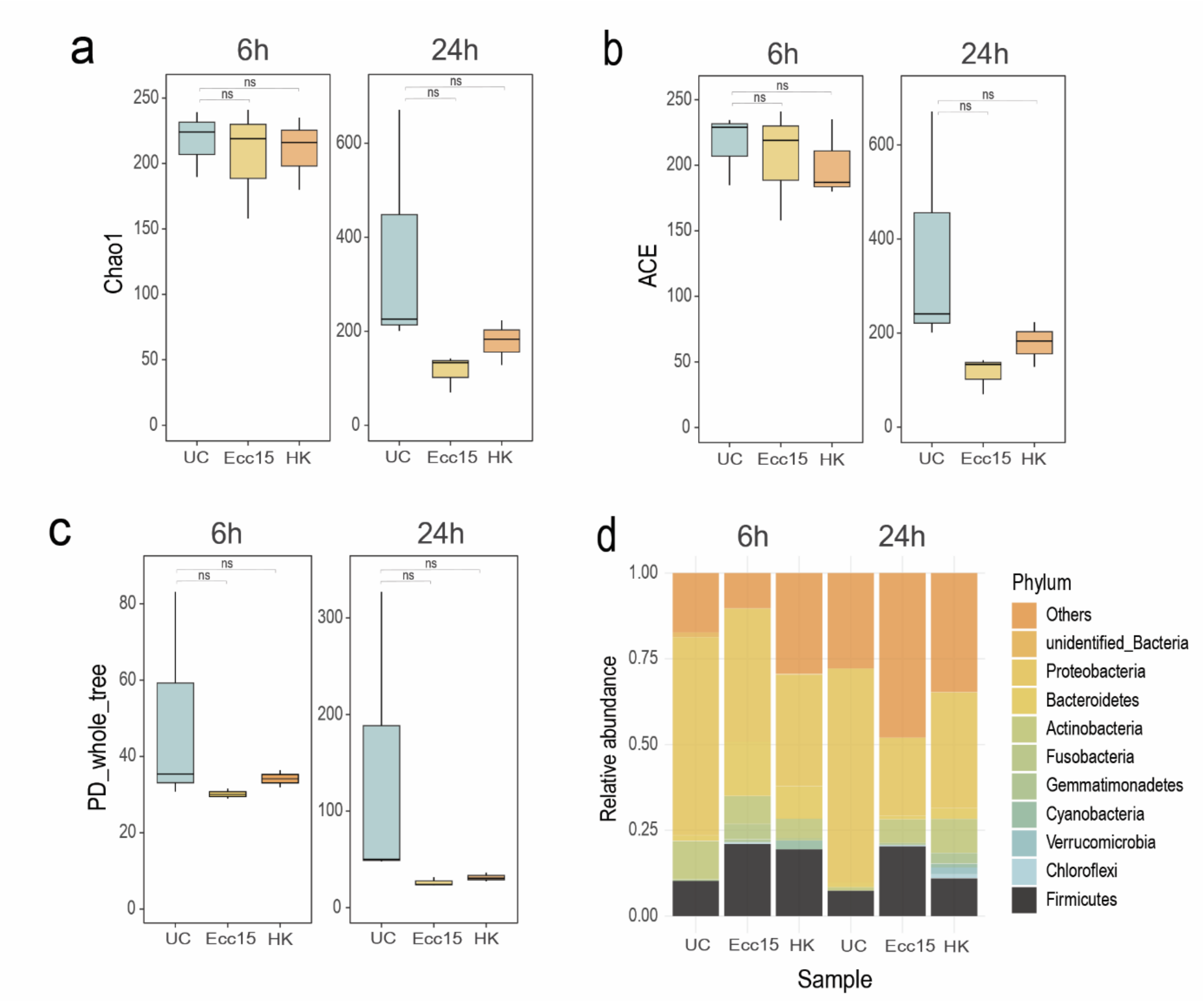


**Supplementary Figure 1. Abundance of *Drosophila* microbiota communities during infection. a-c**, Alpha diversity illustrated by Chao1 (**a**), ACE (**b**), and PD whole tree indexes (**c**), at the 6h and 24h time points after the infection with *Ecc15* alive and heat-inactivated in 10d old conventional files (n=3 independent experiments with 20 guts per treatment and time point). **d**, the relative abundance of 10 dominant OTUs after *Ecc15* infection at the Phylum level. Dot plots and boxplots show median and interquartile range (IQR), and whiskers show either the lower and upper quartiles or range. *P < 0.05, **P < 0.01, ***P < 0.001, ****P < 0.0001. Kruskal–Wallis and Bonferroni post hoc tests were used for statistical analysis.

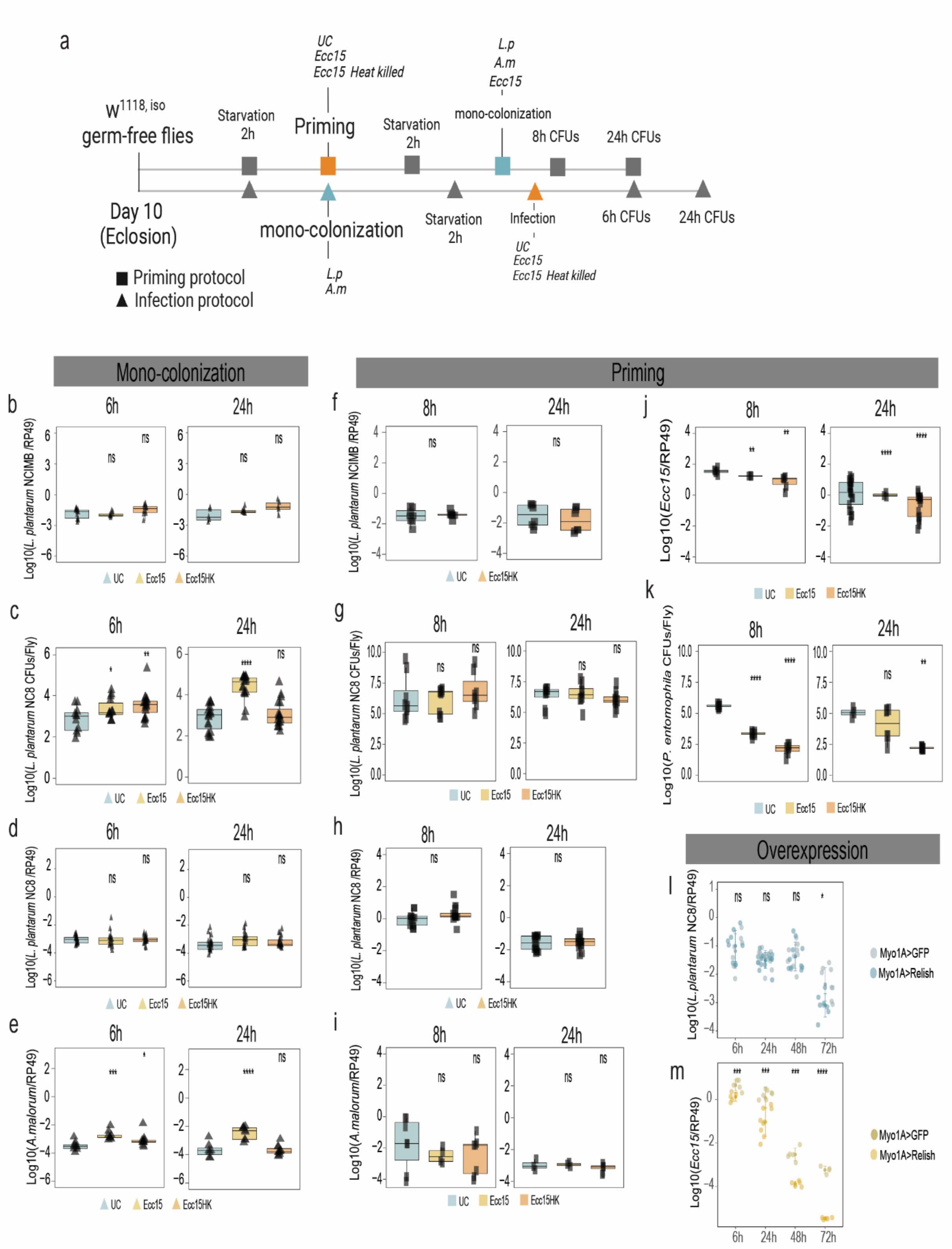


**Supplementary Figure 2. Status of single *Drosophila* microbiota members and pathogens during infection, priming, and overexpression of IMD pathway in *Drosophila* gut. a,** experimental design for Mono-colonization-Infection and Priming protocols. Experiments were performed in 10d old germ-free flies. **Mono-colonization-Infection:** **b**, *L. plantarum*$^{NCIMB}$ loads measured by qPCR after the infection with *Ecc15*. **c-d**, *L. plantarum*$^{NC8}$ loads measured by CFUs (**c**) and qPCR (**d**) at 6h and 24h after the infection with *Ecc15*. **e**, *A. malorum* loads measured by qPCR after the infection with *Ecc15*. (b-e, n=8 independent samples per treatment with 5 flies per sample). The single triangles are mean bacterial load values from pools of n = 5 animals in the Log10 scale. **Priming:** **f**, *L. plantarum*$^{NCIMB}$ loads measured by qPCR after the priming with *Ecc15*. **g-h**, *L. plantarum*$^{NC8}$ loads measured by CFUs (**g**) and qPCR (**h**) at 6h and 24h after the priming with *Ecc15*. **i**, *A. malorum* load measured by qPCR after the priming with *Ecc15*. **j**, *Ecc15* loads measured by qPCR after the priming with *Ecc15*, **k**, *P. entomophila* loads measured by CFUs after the priming with *Ecc1*5. (f-k, n=8 independent samples per treatment with 5 flies per sample). The single squares are mean bacterial load values from pools of n = 5 animals in the Log10 scale. **l-m**, *L. plantarum*$^{NC8}$ (**l**) and *Ecc15* (**m**) loads in *Myo1A-GAL4>UAS-GFP* and *Myo-GAL4>UAS-Relish* flies at 6, 24, 48, and 72h after colonization or infection (n=10 per time point and genotype). Individual dots show mean bacterial load values measured by qPCR from pools of n = 5 animals in the Log10 scale. Boxplots and dot plots show median and interquartile ranges (IQR); whiskers show either lower or upper quartiles or ranges. *P < 0.05, **P < 0.01, ***P < 0.001, ****P < 0.0001. Kruskal–Wallis and Bonferroni post hoc tests were used for statistical analysis.

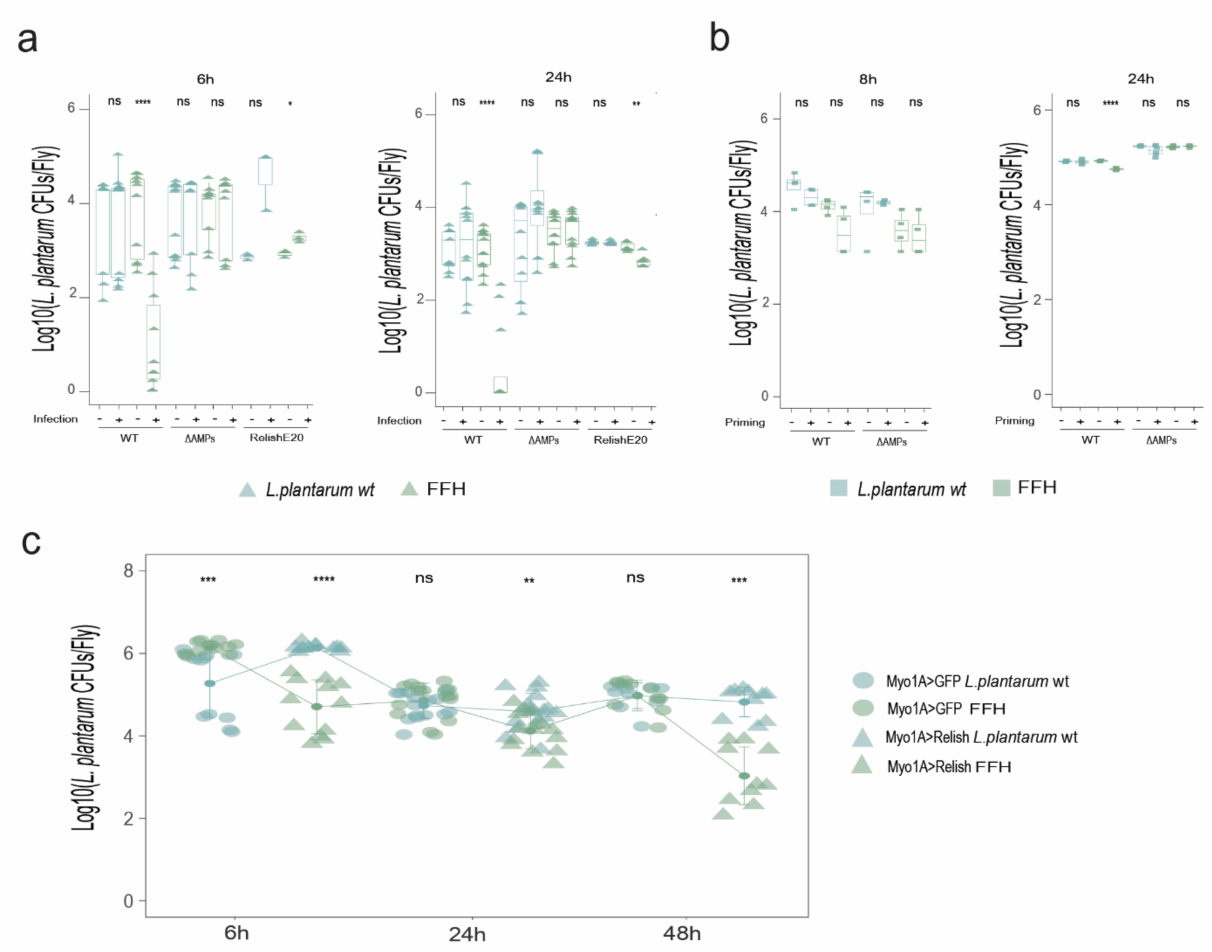


**Supplementary Figure 3.** ***L. plantarum ffh*** **gene is essential to colonize and persist in the gut during infection. a**, *L. plantarum*$^{NCIMB}$ and *L. plantarum*$^{ffh}$ loads in wild type (n=10 per treatment), Δ*AMPs* (n=10 per treatment) and *Relish*$^{E20}$ (n=3 per treatment) flies after the infection with *Ecc15* at 6h and 24h. **b**, *L. plantarum*$^{NCIMB}$ and *L. plantarum*$^{ffh}$ loads after the priming with *Ecc15* at 8h and 24h, in wild type and Δ*AMPs* (n=9 per treatment). **c**, *L. plantarum*$^{NCIMB}$ and *L. plantarum*$^{ffh}$ loads in *Myo1A-GAL4>UAS-GFP* and *Myo-GAL4>UAS-Relish* at 6h (n=12 per treatment), 24h (n=16 per treatment), and 48h (n=10 per treatment) after colonization. Individual tringles, squares and dots show mean CFU values from pools of n = 5 animals in the Log10 scale. Boxplots and dot plots show median and interquartile ranges (IQR); whiskers show either lower or upper quartiles or ranges. *P < 0.05, **P < 0.01, ***P < 0.001, ****P < 0.0001. Kruskal–Wallis and Bonferroni post hoc tests were used for statistical analysis.

**Supplementary tables**

**Supplementary table 2.** Bacterial strains and plasmids used in this study.

| Strain or plasmid | Relevant characteristics | Reference or source |
|---|---|---|
| **Strains** | | |
| *Pectobacterium carotovorum carotovorum (Ecc15)* | Natural *Drosophila* pathogen | 1 |
| *Pseudomonas entomophila* | Lethal intestinal pathogen of *Drosophila* | 2 |
| *Pseudomonas aeruginosa* PA14 | Pathogen used for *Drosophila* intestinal infection | 3 |
| *Levilactobacillus brevis* DSM20556 | | German Collection of Microorganisms and Cell Cultures (DSMZ) |
| *Acetobacter pomorum* WJL | *Drosophila* isolate | 4 |
| *Acetobacter malorum* DSM14337 | | German Collection of Microorganisms and Cell Cultures (DSMZ) |
| *Sphingomonas leidyi* | *Drosophila* isolate | This study |
| *Variovorax* spp | *Drosophila* isolate | This study |
| *Gluconobacter morbifer* G707 | *Drosophila* isolate | 5 |
| *Entorococcus faecalis* | *Drosophila* commensal | 6 |
| *Lactiplantibacillus plantarum* NCIMB 8826 (WCFS1) | Strain with high transformation efficiency | 7 |
| *L. plantarum* NC8 | | 8 |
| *L. plantarum* WJL | *Drosophila* isolate | 5 |
| *L. plantarum* WJL-mCherry | | 9 |
| Δ*dltop* | NC8 strain deleted for *pbpX2* and *dltXABCD* genes | 8 |
| *L. plantarum* P17E3 | NCIMB strain with transposon insertion in *dlt* operon | This study |
| *L. plantarum* P21C9 | NCIMB strain with transposon insertion in *ffh* gene | This study |
| *L. plantarum* NCIMBpVI129 | NCIMB strain carrying pVI129 | This study |
| *L. plantarum* NZ7100 | Derivative of WCFS1 | 10 |

| | | |
|---|---|---|
| *L. plantarum* OatA$^-$ | NZ7100 oatA ::lox72, strain lacking *OatA* | 10 |
| *L. plantarum* OatB$^-$ | NZ7100 oatB ::lox72, strain lacking *OatB* | 10 |
| *L. plantarum* OatAB$^-$ | Strain lacking *OatA* and *OatB* | 10 |
| *L. plantarum* P21C9pGIEB003 | P21C9 mutant containing pGIEB003 plasmid for overexpression of OatA | This study |
| *L. plantarum* P21C9 pNZ8048::ffh | P21C9 mutant containing pNZ8048::ffh plasmid for ffh overexpression | This study |
| *E. coli* TG1 | supE hsd5h thi (Δlac-proAB) F' (traD36 proAB-lacZΔM15) | 11 |
| **Plasmids** | | |
| pVI129 | Ap$^r$ Cm$^r$, pVI1056 containing Phlb A-IS1223ΔIR | 12 |
| pVI110 | Em$^r$, pBR322ori, Pjunc | 12 |
| pNZ8048 | Cm$^r$; shuttle vector containing PnisA promoter and start codon in NcoI site | 13 |
| pGIEB003 | Cm$^r$; pNZ8048 derivative containing *oatA* gene in transcriptional fusion | 10 |
| pNZ8048::ffh | Cm$^r$; pNZ8048 derivative containing *ffh* gene | This study |

**Supplementary table 3.** Primers used in this study.

| **Primer name** | **Sequence 5′→3′** | **Source** |
|---|---|---|
| IRR6 | TCACCGTCATCACCGAAACG | 12 |
| IRL6 | GCCGCACTAGTGATTAAAATAC | 12 |
| *L. plantarum* Fwd | TGGAAACAGATGCTAATACCG | This study |
| *L. plantarum* Rev | GTCCATTGTGGAAGATTCCC | This study |
| *A. malorum* Fwd | TTGACCTTAAGCCGGTGAGC | This study |
| *A. malorum* Rev | TCCCCTACGGCTACCTTGTT | This study |
| *Ecc15* Fwd | GCCGATAGCCCATCAACAGA | This study |
| *Ecc15* Rev | CCTTATGCCTATCGGAGGCG | This study |
| *RP49F* | GACGCTTCAAGGGACAGTATCTG | 14 |
| *RP49R* | AAACGCGGTTCTGCATGAG | 14 |
| *DptA* F | GCTGCGCAATCGCTTCTACT | 14 |
| *DptA* R | TGGTGGAGTGGGCTTCATG | 14 |
| *Socs36E* F | CACAGCAGCAAGCCAGTTTC | This study |
| *Socs36E* R | AGTGCTTTACTGCTGCGACT | This study |
| *Upd3* F | GCGGGGAGGATGTACC | 14 |
| *Upd3* R | GTCTTCATGGAATGAGCC | 14 |
| *Pink1* F | ACGAAATCTTTGGCAACCGC | This study |
| *Pink1* R | TTCCTCAGCGAAAGCGTCAT | This study |
